# *In Vitro* Evaluation of Anti-Inflammatory, Anti-Plaque Efficacy, and Biocompatibility of Norway Spruce (*Picea abies*) Resin Extract for Oral Care Applications

**DOI:** 10.1101/2024.09.17.613464

**Authors:** Kamilla Yamileva, Simone Parrotta, Evgen Multia

**Affiliations:** Faculty of Pharmacy, University of Helsinki, Viikinkaari 5E, 00790 Helsinki, Finland; Repolar Pharmaceuticals Ltd., Nihtisillantie 3, 02630 Espoo, Finland

**Keywords:** spruce resin extract, pinoresinol, gingivitis, periodontitis, anti-inflammatory, antimicrobial, oral health

## Abstract

The periodontal disease is globally highly prevalent, and calls for novel, effective, and preferably bio-based raw materials. Accumulation of dental plaque causes gingivitis, which is reversible by treatments that control the bacterial biofilm. If left untreated, the gingivitis can lead to gingival inflammation and potentially progress to periodontitis. In this study, a natural antimicrobial and anti-inflammatory Norway spruce (*Picea abies*) resin extract was evaluated as a potential option in supportive periodontal care. Lipopolysaccharide-induced macrophage-like cells were used to study the anti-inflammatory properties *in vitro*. The spruce resin extract at 20% concentration had the highest anti-inflammatory effect, comparable to a corticosteroid’s effect on pro-inflammatory cytokines interleukin-1 beta (IL-1β), tumor necrosis factor-alpha (TNF-α), and matrix metalloproteinase-3 (MMP-3). Consequently, the 20% spruce resin extract was selected for toothpaste formulation. Its anti-plaque efficacy was evaluated by total aerobic colony counts and the proportions of streptococci grown on the surfaces of the treated glass rods using pooled human saliva. It was found that the toothpaste effectively reduced dental plaque biofilm, matching the anti-plaque efficacy of Corsodyl mouthwash, containing chlorhexidine digluconate. The toothpaste was also found to be non-corrosive in biocompatibility studies on three-dimensional (3D) models of human oral and gingival epithelium. These findings provide scientific validation of spruce resin’s effectiveness in oral care, elucidating probable reasons why people have historically chewed resins for oral care purposes.

**Highlights:**

- Spruce resin extract was studied for oral care applications.
- It had anti-inflammatory effect on cytokines IL-1β, TNF-α, and MMP-3.
- 20% spruce resin extract toothpaste was developed.
- The spruce resin extract toothpaste effectively reduced dental plaque biofilm.
- The toothpaste was non-corrosive on 3D models of oral and gingival epithelium.

**Graphical abstract:** 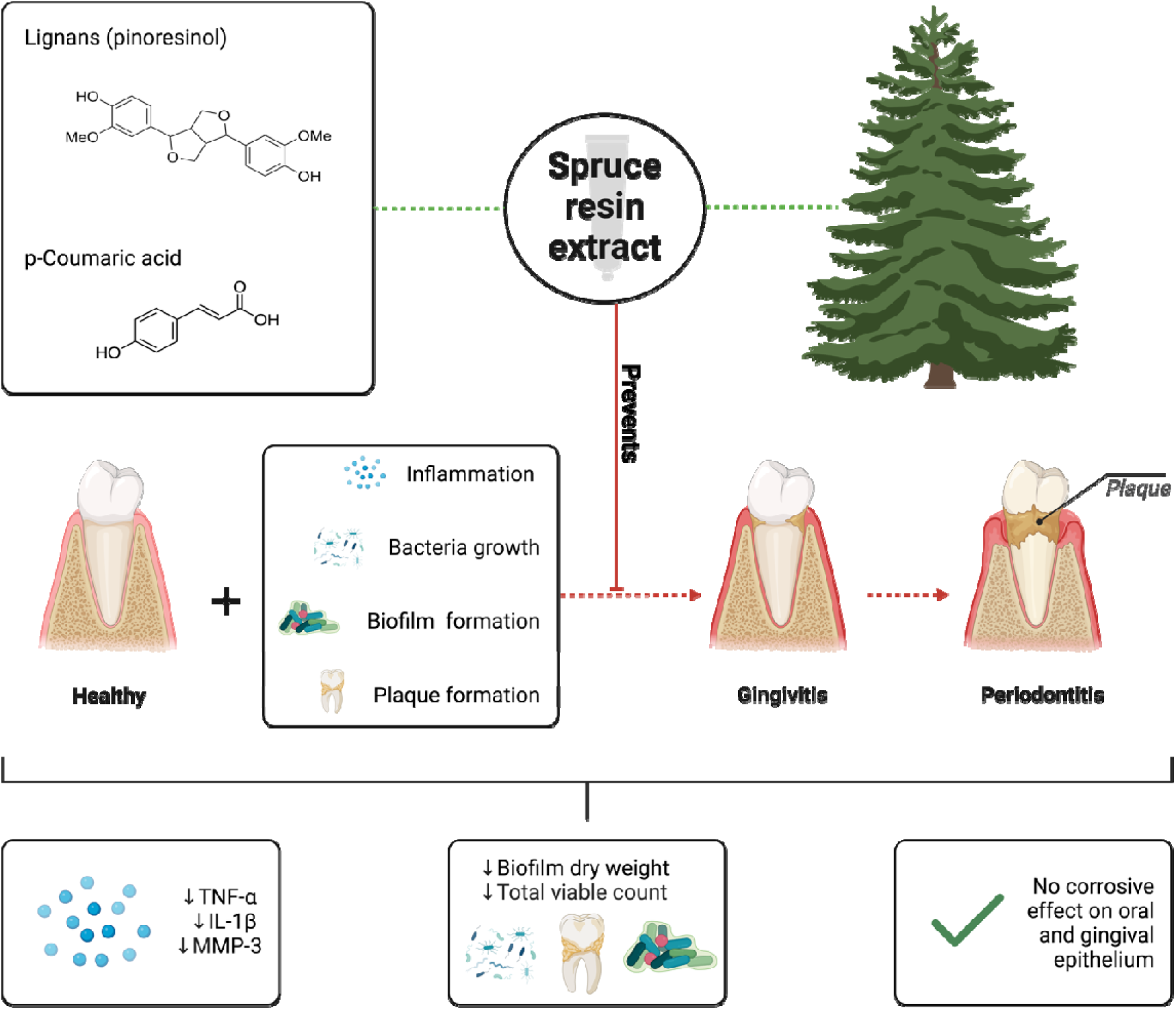

## 1. Introduction

Dental plaque (a whitish bacterial biofilm) accumulates on teeth and gums, particularly around the gum line [1]. Overall, the oral microbiome contains more than 700 bacterial species [2]. *Streptococcus mutans* is the main cause of dental plaque formation that leads to inflammatory response in the oral and gingival epithelium [3,4]. An increased amount of plaque at the gingival margin causes an inflammatory response that leads to increased gingival redness, swelling, and bleeding [1]. This is because pathogenic bacteria trigger the innate immune system to release pro-inflammatory mediators, such as IL-1β, TNF-α, and MMP-3 [5–12]. The increase of pathogenic bacteria in the oral cavity tends to replace beneficial bacteria, leading to an imbalance that favors the growth of pathogenic bacteria, and results in additional plaque accumulation [13].

By maintaining good oral hygiene, it is possible to prevent the formation of dental cavities, halitosis, and periodontal disease (at early-stage gingivitis and later periodontitis) [1,5]. The conventional treatment options for dental plaque care include mechanical removal [14] and the use of antimicrobial agents (e.g., chlorhexidine) [15]. However, these might cause irritation and imbalances in oral microbiota [13,16]. Thus, there is a need for biocompatible formulations preferably based on natural compounds targeting oral pathogens and their biofilms [17].

Natural compounds are an underexplored source of safe, effective, and sustainable antimicrobials for supportive periodontal care [18]. The main benefit of plant-based antimicrobials is that they do not usually have the side effects associated with synthetic chemicals and have limited reports of antimicrobial resistance development. For example, natural resins have traditionally been used in medicine to treat acute and chronic infected wounds, sores, pressure ulcers, punctured abscesses, suppurating burns, onychomycosis, paronychia, and ear infections [19,20]. Tree resins have also been used as chewing gums for teeth cleansing and breath freshening by the ancient Egyptians, the Mayan Indians, and the American Indians [21,22]. Chewing resinous gums has been a common practice to promote oral hygiene and freshen the breath, as these natural gums not only help to clean the teeth mechanically but also provide antimicrobial benefits due to the resin’s bioactive compounds [23]. One of the promising natural raw materials for oral care applications is coniferous spruce (*Picea abies)* resin and its water-based extract [24] due to their anti-inflammatory and antimicrobial properties [25–32].

In this study, *in vitro* anti-inflammatory properties of spruce resin extract and pinoresinol (lignan found at high concentrations in spruce resin extract) were evaluated with U937 macrophage-like cells. In addition, a toothpaste formulation containing 20% spruce resin extract was developed and tested for its anti-plaque and antimicrobial efficacy against streptococci. Finally, the toothpaste’s biocompatibility was evaluated in human gingival and oral epithelium 3D cell models. The objective of this study was to investigate the potential of spruce resin extract as a natural and biocompatible agent for supportive periodontal care.

## 2. Methods

### 2.1. Study 1: Effects of Spruce Resin Extract and Pinoresinol on LPS Induced IL-1β, TNF-α, and MMP-3 Production in U937 cells

#### 2.1.1. Preparation of Spruce (*Picea abies*) Resin Extract

The spruce resin extract was prepared according to a patented method [24]. In brief, spruce resin was mixed in ratio 1:9 (w/w) in glycerol (Sigma-Aldrich, St. Louis, USA). Followed by dilution of glycerol-spruce resin dispersion further with a physiological saline solution (Sigma-Aldrich, St. Louis, USA), in a w/w ratio of 1:1. Resulting spruce resin extract was filtered through 1 µm filter before further use.

#### 2.1.2. Characterization of Spruce Resin Extract by Gas Chromatography Mass Spectroscopy (GC-MS)

0.5 mL of the spruce resin extract solution was extracted with 2 mL of dichloromethane (containing 100 ppm of heptadecanoic acid and betulin) (Sigma-Aldrich, St. Louis, USA). Additional 2 x 2 mL of dichloromethane extractions were done with vortexing. Dichloromethane (Sigma-Aldrich, St. Louis, USA) was evaporated and silylation was done with 0.5 mL of TMSI (Sigma-Aldrich, St. Louis, USA).

Silylated samples were analyzed with GC-MS (Shimadzu GCMS-QP2020SE, Japan) and helium was used as carrier gas. Column was HP-5 (length 30 m x 0.32 mm ID x 0.25 µm df, Agilent, USA). The chromatographic conditions were following: initial temperature 170 °C; temperature rate 5 °C/min; final temperature 300 °C for 25 minutes; injector temperature 300 °C and split ratio 1:10. Identifications of the extractives were done by comparison of the GC retention time and the EI MS spectra with compounds characteristic m/z found in the literature and publicly available databases (NIST14.L). The extractions and analysis were done in 3 technical repeats.

#### 2.1.3. Preparation of Pinoresinol solution

A saturated stock of pinoresinol (Sigma-Aldrich, St. Louis, USA) was prepared by aseptically adding the pinoresinol 1%, 10% or 20% (v/v) to 10 mL of Vehicle (25% Glycerine/dH_2_O) and warmed to 37°C for 10 minutes with gentle mixing. This solution was then mixed gently at room temperature for 2.5 hours before filtering through a 2.5 µm filter.

#### 2.1.4. Preparation of Controls for U937 experiments

A stock of dexamethasone (Sigma-Aldrich, St. Louis, USA) was prepared aseptically in ethanol (VWR Chemicals, Fontenay-sous-Bois, France) as a positive control. This was further diluted in complete media (CM) to give a 2X working stock (1 µM) and stored at 4-8°C for the duration of the study. The final concentration in the well was 0.5 µM. Vehicle was 25% of Glycerine (Sigma-Aldrich, St. Louis, USA) in distilled water.

#### 2.1.5. Preparation of Complete media (CM)

To formulate complete media, RPMI-1640 (IM) (Invitrogen, Massachussets, USA) was supplemented with 10% heat inactivated fetal bovine serum (FBS) (Invitrogen, Massachussets, USA), and 100 U/ml penicillin and 100 µg/ml streptomycin (Invitrogen, Massachussets, USA). Complete media (CM) was prepared aseptically and stored at 4-8°C for the duration of the study. Complete media was warmed to 37°C immediately prior to use.

#### 2.1.6. Preparation of Phorbol 12-myristate 13-acetate (PMA)

A 10X working stock (2000 nM) of phorbol 12-myristate 13-acetate (PMA) (Sigma-Aldrich, St. Louis, MO) was prepared aseptically in CM and stored at 4-8°C for the duration of the study. The final concentration in the well was 200 nM.

#### 2.1.7. Preparation of Lipopolysaccharide (LPS)

A 10X working stock (10 ug/mL) of LPS from *S. abortus equi* (Enzo Life Sciences, Farmingdale, USA) was prepared aseptically in IM and stored at 4-8°C for the duration of the study. The final concentration in the well was 1 µg/mL.

#### 2.1.8. Preparation of Growth Media

RPMI-1640 with 4.5 g/L glucose with L-glutamine (GlutaMAX) +10% heat inactivated FBS +100 U/mL penicillin + 100 µg/mL streptomycin (complete medium: CM).

### 2.2. Experimental Flow of Study 1

#### 2.2.1. Effects of Spruce Resin Extract and Pinoresinol on LPS induced IL-1β, MMP-3 and TNF-α Production in U937 cells

Cells from U937 human monocyte cell line (HPA Culture Collection, 85011440), were treated with 200 nM PMA for 72 hours to differentiate them into macrophages. PMA was then removed, and the cells incubated with media control, vehicle control, 0.5 µM dexamethasone or 2 test items: 1%, 10% or 20% (v/v) spruce resin extract, or 1%, 10% or 20% (v/v) pinoresinol for 1 hour. After 1 hour the cells were treated with 1 µg/mL of LPS, unstimulated cells were treated with incomplete media of the same volume.

After 20 hours of LPS stimulation cell viability was assayed by adding AlamarBlue™ (Invitrogen, Massachussets, USA) to each well, incubating for 1 hour and measuring the fluorescence (544 nm excitation and 590 nm emission).

After 24 hours of LPS stimulation the well supernatants were harvested and subsequently analyzed for IL-1β, MMP-3 and TNF-α (Procarta Multiplex Assay, Thermo Fisher Scientific, USA) levels by Luminex multiplex according to manufactures’ instructions.

#### 2.2.2. Schematic Depiction of *in vitro* Stimulation of U973 cells

Schematic Depiction of *in vitro* Stimulation of U973 cells can be found in Figure 1.

**Figure 1.**
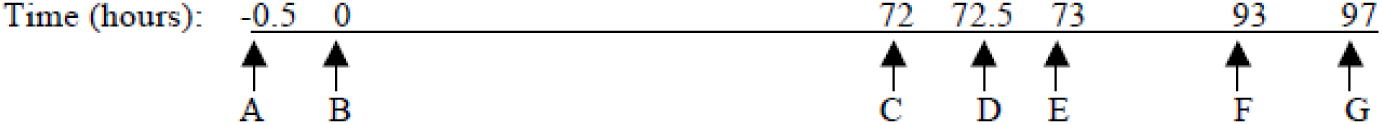
Schematic depiction of in vitro Stimulation of U973 cells. (A)= Plate U937 cells, (B)= Add PMA, (C)= Replace media, (D)= Add test items and reference compounds, (E)= Add LPS, (F)= AlamarBlue added, (G)= Cells assayed for viability. Harvest cell culture supernatants for cytokine analysis. Cells were allowed to settle and incubated at 37°C with 5% CO_2_.

##### 2.2.2.1. (A) Cell Culture Set Up

Cells were thawed, washed in CM and repeatedly passaged before use. A solution containing 5x10^5^ cell/mL in complete media was prepared. 5x10^4^ cells were added in 0.1 mL to the appropriate wells of 96 well black walled plates. Cells were allowed to settle for 30 minutes.

##### 2.2.2.2. (B) PMA addition

A stock solution of PMA was prepared (2000 nM). This was diluted in the well to give a final PMA concentration in the well of 200 nM. An equal volume of complete media was added to the non-PMA treated cells. Cells were incubated for a further 72 hours.

##### 2.2.2.3. (D) Treatment addition

After PMA treatment, the media was aspirated from wells and replaced with complete media. Test items, reference articles or vehicle were then added to the appropriate wells. An equal volume of complete media was added to the non-treated wells. Following control or test item addition to the wells, the plates were incubated at for 1 hour.

##### 2.2.2.4. (E) LPS stimulation

A stock solution of LPS was prepared (1000 µg/mL). This was further diluted in the well to give a final LPS concentration in the well of 1 µg/mL. An equal volume of incomplete media was added to the non-stimulated wells. Cells were incubated for a further 20 hours.

##### 2.2.2.5. (F) Addition of AlamarBlue™

After 20 hours of LPS stimulation AlamarBlue™ was added to each well. The plates were incubated for a further 4 hours.

##### 2.2.2.6. (G) Culture termination, Sample Collection, and Cytokine/Chemokine Statistical Analysis

At 24 hours post LPS stimulation cell viability was assayed by reading the fluorescence of AlamarBlue™, 544 nm excitation and 590 nm emission. Supernatants were then harvested and stored at -80°C until assayed.

Cell culture supernatants were assayed for IL-1β, MMP-3, and TNF-α using a Luminex-based assay according to the manufacturer’s instructions. Data were collected using a Luminex 100 (Luminex Corporation, Austin, TX). Standard curves were generated using a 5-parameter logistic curve-fitting equation weighted by 1/y (StarStation V 2.0; Applied Cytometry Systems, Sacramento, CA). Each sample reading was interpolated from the appropriate standard curve.

The mean and the standard error of the mean was calculated for all treatments assayed in triplicate. The significance of treatment effect was determined by ANOVA with Tukey post-hoc analysis, a p-value of ≤ 0.05 was considered significant.

### 2.3. Study 2: *In vitro* Study to Compare the Anti-Plaque Efficacies of Spruce Resin Toothpaste Formulation Versus Enzymatic Toothpaste, and a Positive and Negative Control

In this *in vitro* method [33], the resin extract toothpaste formulation was compared to enzymatic toothpaste and controls in its efficacy to prevent plaque biofilms from growing onto the surfaces of roughened glass rods.

Dental plaque was grown over 3 days by immersing the roughened glass rods into fresh, pooled human saliva containing 0.1% sucrose. During treatment days 2 and 3, plaque growth was also encouraged by periodically exposing the glass rods to a nutrient broth containing tryptone soya broth (TSB), saliva, and sucrose (Sigma-Aldrich, St. Louis, USA).

On treatment day 1, the roughened glass rods were treated with a single application of the assigned treatment. On treatment days 2 and 3, the roughened glass rods were treated with two applications of the assigned treatment.

At the end of the 3 days, the plaque biofilms were harvested from the roughened glass rods, and the level of plaque growing on the roughened glass rods was quantified by dry weight (g) and the total viable bacterial counts (TVC). Lower dry weights and TVC counts were indicative of greater anti-plaque efficacy. The proportions of streptococci within the total aerobic counts (TVC) were also evaluated.

#### 2.3.1. Glass Rod Substrate

Glass rods (6 mm in diameter and 10 cm in length) were used as a substrate for plaque growth, with 8 rods assigned to each treatment group. All glass rods were roughened to a consistent finish along two-thirds of their length with 400-grit silicon carbide paper. The rods were then rinsed thoroughly with deionized water before being air-dried. Finally, the rods were sonicated to remove loose debris and sterilized by immersion in 70% isopropanol (Sigma-Aldrich, St. Louis, USA) for 1 hour. This step was performed inside a microbiological safety cabinet (Bassaire, UK).

#### 2.3.2. Substrate Holder

A 6 mm diameter hole for each rod was drilled into the center of Falcon tube lids. These lids were sterilized in 70% ethanol for 1 hour. This step was performed inside a microbiological safety cabinet. Each Falcon tube housed one of the 8 rods assigned to each treatment group.

#### 2.3.3. Saliva Containing 0.1% Sucrose

All saliva was collected in accordance with Intertek CRS standard operating procedures. Written informed consent was provided by Intertek CRS staff donors, who provided stimulated saliva (using flavorless gum) into 30 mL containers. All anonymized saliva donations were recorded using a donor log, with fresh saliva collected and pooled during each study day. When a sufficient volume of saliva was collected, 0.1% sucrose (w/w) was added.

#### 2.3.4. Nutrient Broth

The nutrient broth consisted of a 3% tryptone soya broth (TSB) solution, containing 10% sucrose (w/w), and 6.5% fresh, pooled human saliva (w/w). The nutrient broth containing 10% sucrose was made in advance and sterilized by autoclaving. Before performing the nutrient immersions, 6.5% of fresh, pooled saliva (w/w) was added to the nutrient broth.

#### 2.3.5. Preparation of Spruce Resin Extract Toothpaste

20 % spruce resin extract (containing water, glycerol, spruce resin extractives, and sodium chloride) was used to prepare a spruce resin extract toothpaste. Other ingredients (Sigma-Aldrich, St. Louis, USA) of the toothpaste were hydrated silica, sorbitol, aroma, xanthan gum, papain, bromelain, amyloglycosidase, citric acid, sodium benzoate, and potassium sorbate.

#### 2.3.6. Treatment Preparation

All controls were pipetted on day one and then stored at room temperature. The control products were decanted in 4 mL aliquots into each Falcon tube. All toothpaste formulations were mixed with sterile deionized water 30 minutes before treatment. The proportion was 1 part toothpaste to 1.6 parts sterile deionized water. The toothpaste formulations (Table 1) were homogenized using an end over mixer, and 4 mL aliquots of the toothpaste slurries were decanted into the Falcon tubes.

**Table 1.**
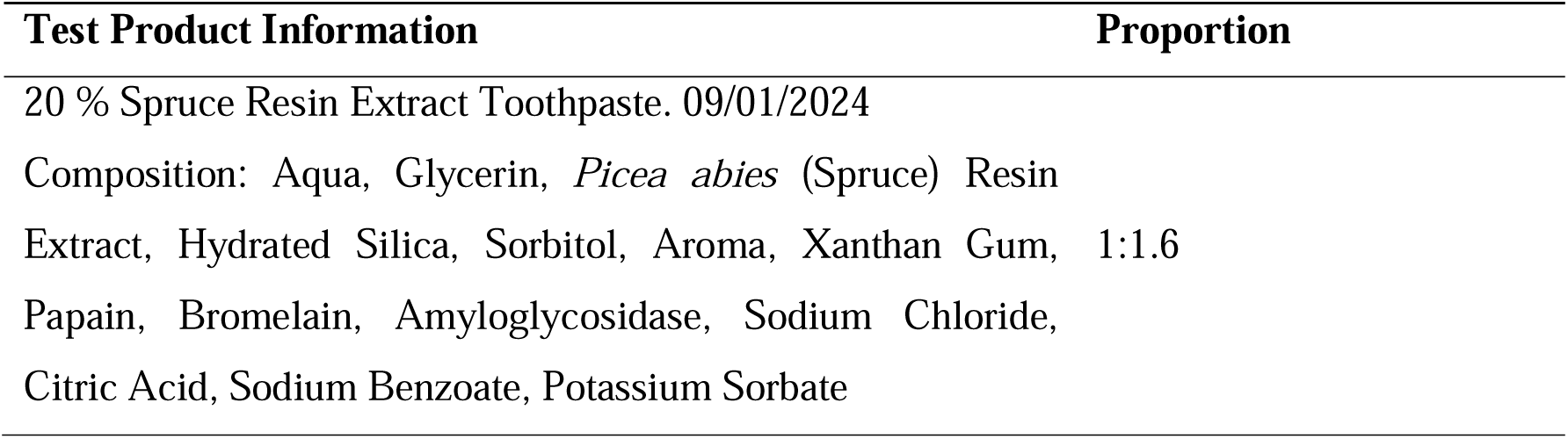
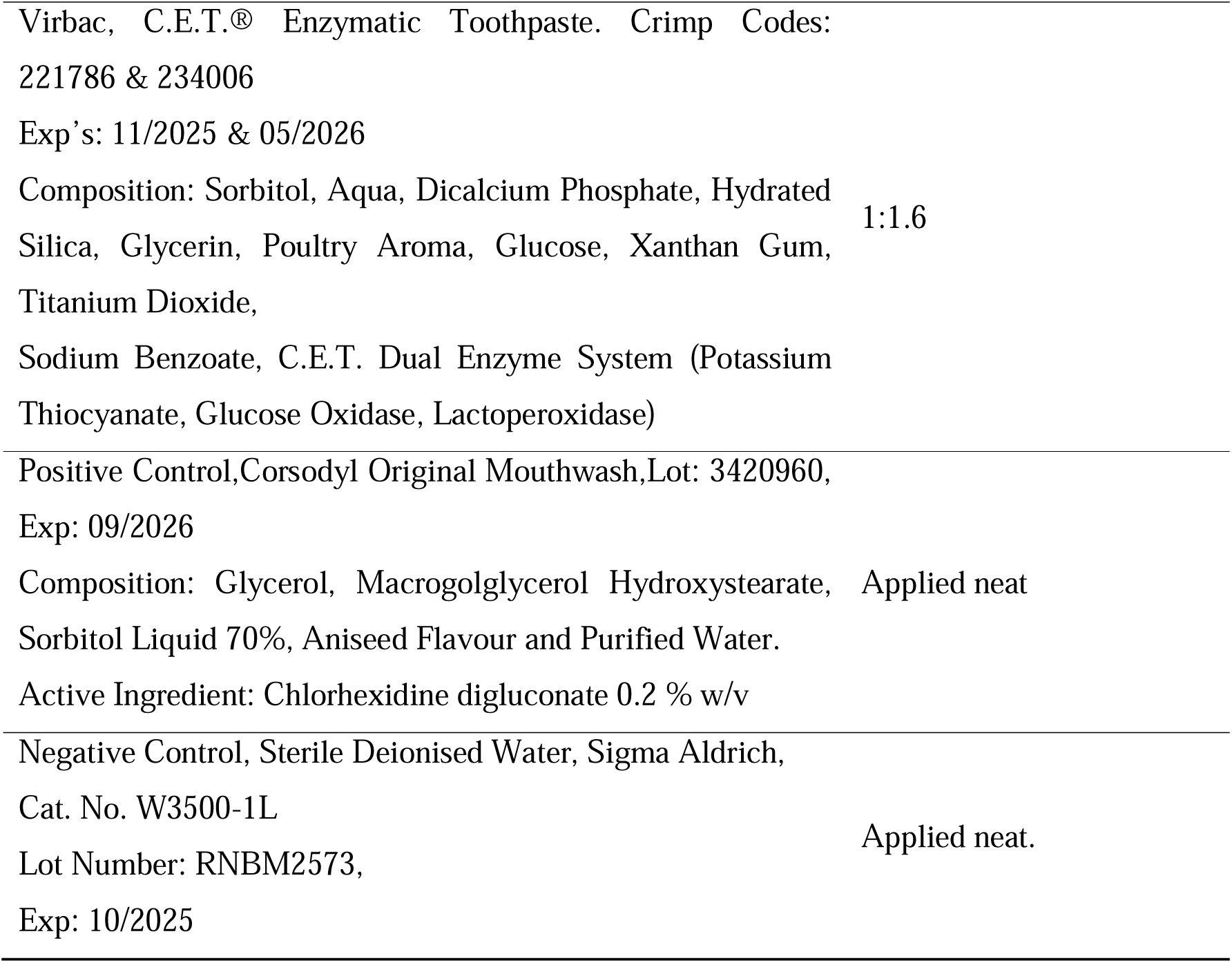
Test products used in the *in vitro* study to compare the anti-plaque efficacies.

#### 2.3.7. Treatment Procedure

The sterilized rods were inserted into the lids ready for treatment inside the microbiology safety cabinet. The rods and lids were used as one from this point onwards.

On the day 1, the sterilized glass rods were pre-treated with 4 mL of their respective treatment. After 2 minutes, the rods were removed, rinsed with sterile water, and then placed into 4 mL of saliva (containing 0.1% sucrose). The tubes were transferred to an incubator (LEEC, Nottingham, UK) for 18 hours at 37°C, whilst being agitated on a plate shaker set to 250 rpm. On the days 2 and 3, the rods were once again treated, before being placed into a nutrient broth and transferred back into the incubator and agitated. After 6 hours in the broth, the rods were treated again, transferred into saliva containing 0.1% sucrose, and placed in the incubator for a further 18 hours. On the day 4, half of the glass rods per group were retained for dry weight analysis. The other half of the rods were subjected for TVC analysis.

#### 2.3.8. Dry Weight Analysis

The assigned four rods were analyzed to determine the dry weight of the biofilms growing on the glass rods. All rods were allowed to air dry. Rods were weighed on an analytical balance. After rehydration, the biofilm was removed, and the rods were reweighed.

#### 2.3.9. Total Aerobic Count Analysis (TVC, Total Viable Counts) and Proportions of streptococci within the Biofilms

Measurement of the total aerobic counts (TVC) was performed with 4 rods per treatment group, The rods were immersed in sterile phosphate-buffered saline (PBS) solution. All rods were then dip-washed 3 times in sterile saline and the original sample pots were rinsed. The sample pots were filled with 20 mL sterile saline, and the rods were scraped until visibly clean. The test material was mixed by vortexing (Stuart) to obtain a homogenous suspension.

The suspension was streaked out onto Columbia blood agar and Potato Dextrose agar. Serial dilutions were also plated out using Tryptone Soy agar to achieve a countable level. The plates were incubated at 37°C (± 2°C) according to standard practice. In addition to determining the TVC, the proportions of streptococci within the total viable count were also evaluated.

#### 2.3.10. Statistical Analysis

Minatab21 was used to generate descriptive statistics and to statistically compare the post-treatment plaque dry biofilm weights and the total aerobic colony counts (TVC). Descriptive statistics were calculated for the plaque dry biofilm weight data. A General Linear Model ANOVA was selected to statistically compare the post-treatment plaque dry biofilm weights and the total viable aerobe counts achieved by the different treatments. A Tukey test was used to make pairwise statistical comparisons between the test products and the controls. A p-value of ≤ 0.05 was considered significant.

### 2.4. Study 3: Oral and Gingival Epithelium Corrosion Tests

An *in vitro* test [34,35] (Normative Reference: OECD Technical Guideline 431) was performed to evaluate the oral and gingival epithelium corrosive effect of the product. An *in vitro* reconstituted human 3D-gingival epithelium model (type Episkin, Lot. No. 24 HGE 008) and 3D-oral epithelium model (type Episkin, Lot. No. 24 HOE 0010) were used. Prior to performing the test procedure, the oral and the gingival epithelium model cell culture inserts were incubated for 24 h at 37°C and 5 % pCO2 in a cell culture incubator in fresh maintenance medium. After this preincubation time 100 µl of PBS was pipetted on the surface of each oral or gingival epithelium model. Then 100 µl glacial acetic acid (Sigma-Aldrich, St. Louis, USA) was pipetted on one group of these oral or gingival epithelium inserts as a (gingival and oral epithelium corrosive) positive control. 100 µl PBS solution was pipetted on a second oral or gingival epithelium insert group as a (non-gingival and non-oral epithelium corrosive) negative control. 100 mg of the spruce resin toothpaste was applied on a third oral or gingival epithelium insert group. Test procedure was performed in two incubation clusters. The first incubation cluster was incubated for 3 min at room temperature. The second cluster was incubated at room temperature for 1 h. After incubation all inserts were rinsed thoroughly with sterile PBS, blotted on a paper towel to remove excess PBS, and an MTT-test was performed to measure the cell vitality of the gingival epithelium inserts.

### 2.5. Measurement of cell vitality (MTT-test) for Oral and Gingival Epithelium Corrosion Tests

The inserts were rinsed once in maintenance medium using a 24-well cell culture plate and transferred in a second 24-well cell culture plate with 300 µl maintenance medium containing 1 mg/mL MTT (Sigma M5655). All inserts were incubated for additional 1.5 h in a cell culture incubator at 37°C and 5 % pCO2. Afterwards all inserts were blotted on a paper towel and the MTT-dye was extracted from the oral or gingival epithelium samples using 0.8 ml isopropanol (Sigma-Aldrich, St. Louis, USA) per insert. The extinction of each isopropanol extract was measured in a photometer at 570 nm to obtain data for the relative vitality of the cells in the oral or the gingival epithelium samples compared to the negative control (PBS).

## 3. Results

### 3.1. Gas Chromatography Mass Spectrometry (GC-MS) Analysis of Spruce Resin Extract

The spruce resin extract was analyzed with GC-MS and the chemical composition is presented in Table 2. Heptadecanoic acid and betulin were used as internal standards for the quantification of the compounds found in the extract. The extract contained mostly p-coumaric acid (414.8±5.5 ppm) and lignans (647.4±9.0 ppm). Pinoresinol had the highest concentration (428.1±8.0 ppm) among lignans. Other lignans found in the extract were isolariciresinol (156.8±1.4 ppm) and secoisolariciresinol (62.5±0.6 ppm). The sum of oxidized resin acids (32.1±1.0) is stated as the minimum amount due to the possibility of some resin acids not eluting from the GC column [28] and the limitations of the derivatization process of resin acids with TMSI, which may result in incomplete detection.

**Table 2.**
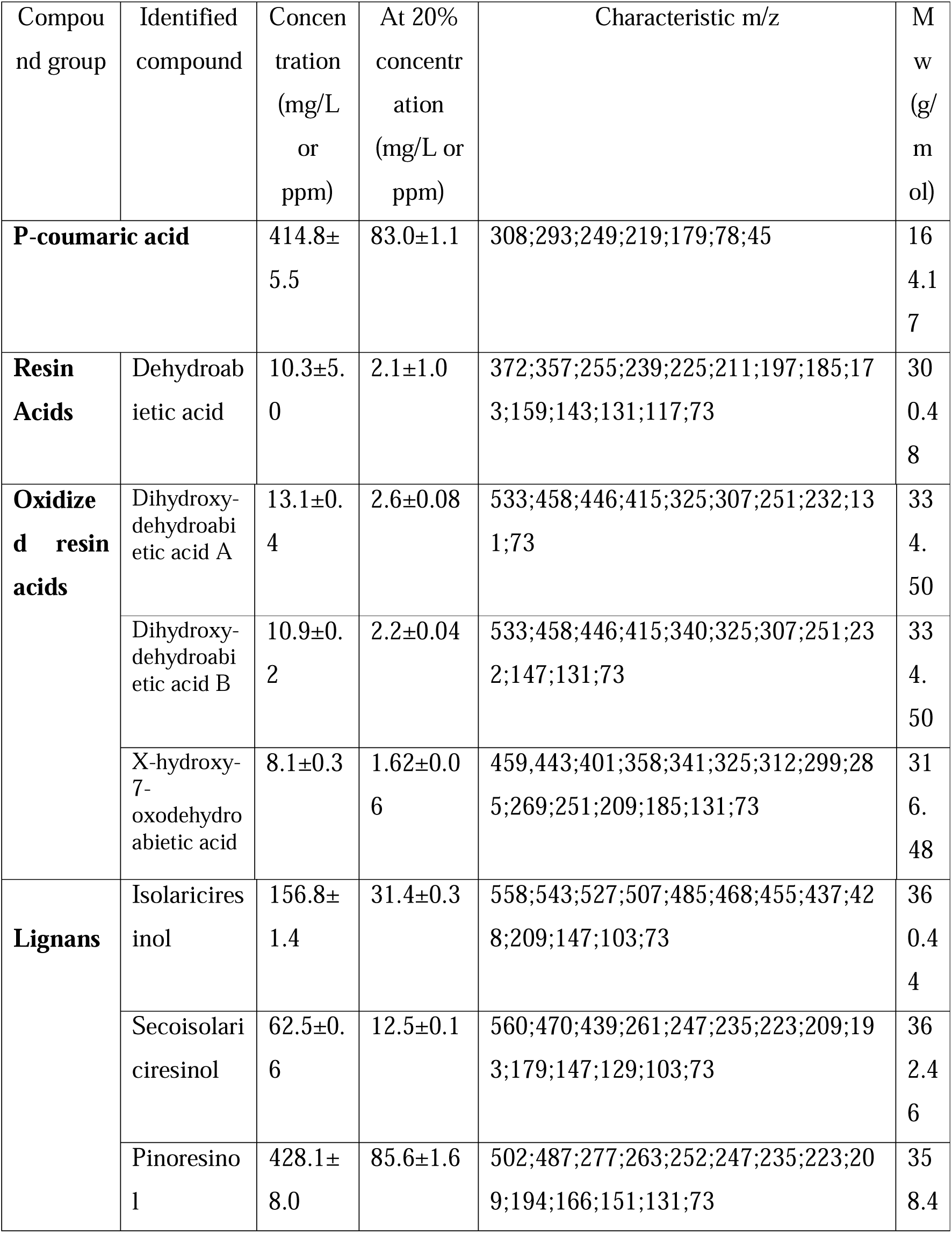
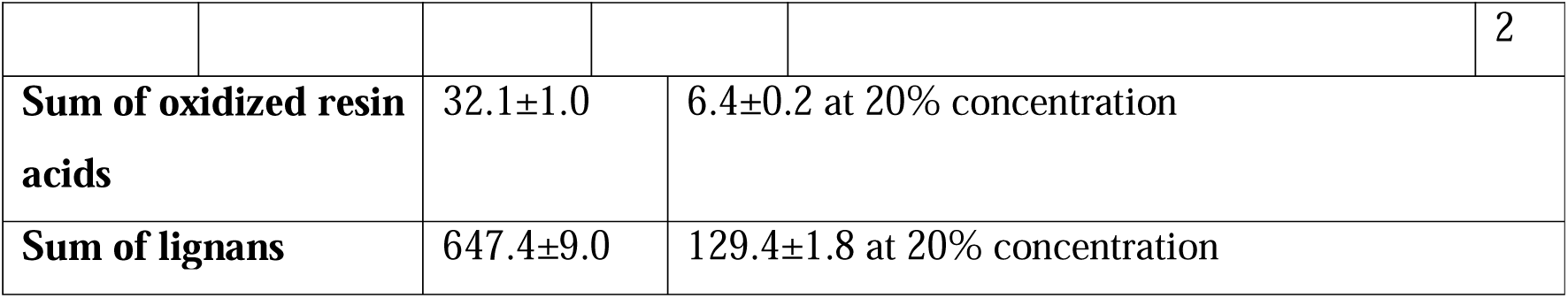
Chemical composition of spruce resin extract analyzed by GC-MS total-ion chromatograms of TMSI silylated compounds. Standard deviations are based on three independent measurements (n=3). MS spectra were referenced with literature and publicly available databases. 20% concentration of the spruce resin extract was used for the toothpaste formulation.

### 3.2. Biocompatibility of Spruce Resin Extract and Pinoresinol **–** U937 Cell viability with AlamarBlue Assay

To study the viability of U937 cells, an AlamarBlue™ assay was used 24 hours after lipopolysaccharide (LPS) addition. The fluorescence intensity (FI) detected from the AlamarBlue™ assay reagent increased with metabolic activity within the culture system, which indicated cell viability. The LPS-stimulation of U937 cells increased significantly their number compared to unstimulated, untreated cells (Figure 2A). A significant decrease in mean AlamarBlue™ FI was observed between PMA-treated, LPS-stimulated cells and either PMA-untreated, LPS-unstimulated cells or PMA-treated, LPS-unstimulated cells. No significant difference in mean AlamarBlue™ FI was observed between PMA-treated, LPS-stimulated cells compared dexamethasone-treated, LPS-stimulated cells. Treatment of LPS-stimulated cells with 0.5 µM dexamethasone, 1% vehicle or 10% vehicle did not affect cell viability. Vehicle was 25% of glycerine in distilled water. However, 20% vehicle significantly reduced cell viability compared to PMA treated, LPS-stimulated cells. Treatment of LPS-stimulated U937 cells with 1% - 20% (v/v) spruce resin extract, or 1% or 10% (v/v) pinoresinol did not significantly affect cell viability compared to their respective vehicle control. Interestingly, at higher concentrations (20%), pinoresinol appears to increase cell viability compared to 20% vehicle treated cells, indicating a potential protective effect or enhancement of cell proliferation. However, the spruce resin extract did not show this effect at any concentrations, even though it contains high proportion of pinoresinol (Table 2).

**Figure 2.**
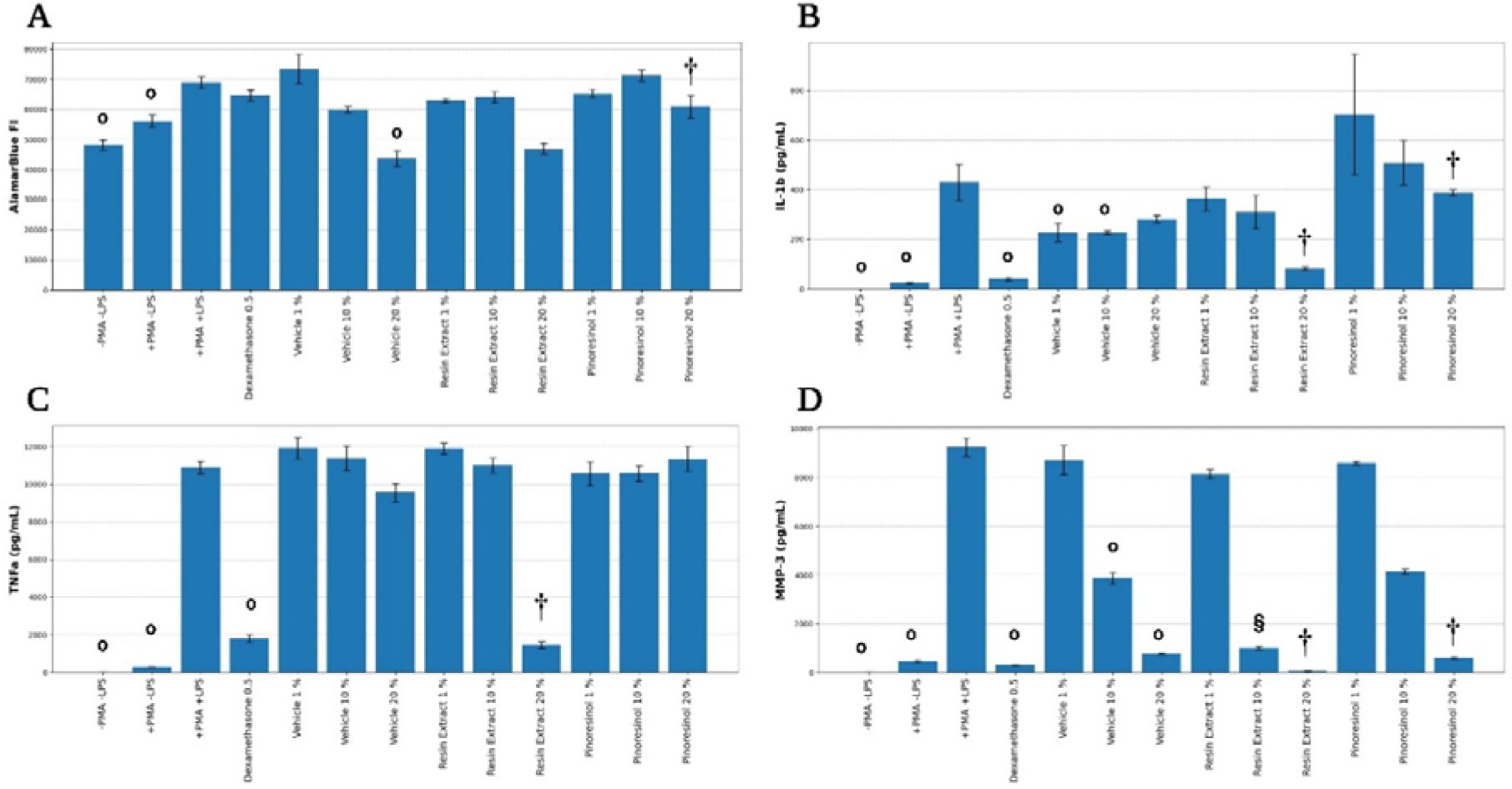
U937 cells. (A) Cell viability by AlamarBlue™ assay. (B) IL-1β production assay. (C) TNF-α production assay. (D) MMP-3 production assay. All assay experiments were done in 3 technical replicates (Mean ± SEM, n=3). Results were statistically compared by ANOVA with Tukey post-hoc analysis. **°** treatment significantly (p ≤ 0.05) different to PMA treated, LPS-stimulated cells. **§** treatment significantly (p ≤ 0.05) different to 10% vehicle in LPS-stimulated cells. **†** Treatment significantly (p ≤ 0.05) different to 20% vehicle in LPS-stimulated cells. PMA= phorbol 12-myristate 13-acetate, LPS= Lipopolysaccharide.

### 3.3. Anti-Inflammatory Action of Spruce Resin Extract **–** Cytokine and Chemokine Levels of U937 Cells

Following the assessment of the cell viability, cell culture supernatants were harvested and analyzed for the presence of IL-1β, TNF-α, MMP-3 by Luminex™ assay. Part of the data was included in a patent EP3247370B1 [24]. LPS-stimulation of U937 cells significantly increased the levels of the cytokines IL-1β and TNF-α, as well as MMP-3, while treatment with dexamethasone (an anti-inflammatory glucocorticoid) significantly reduced the levels of these cytokines (Figure 2B-D). In the LPS-stimulated cells, treatment with 1%, 10%, or 20% vehicle did not significantly affect the levels of TNF-α production (Figure 2C). 1% and 10 % vehicles significantly reduced the level of IL-1β produced, and 10% and 20% vehicles significantly reduced the level of MMP-3 produced. Both IL-1β and TNF-α levels detected within the supernatants of LPS-stimulated U937 cells were significantly reduced following treatment with spruce resin extract at 20% when compared to 20% vehicle-treated cells. A significant increase of IL-1β in 20% pinoresinol treated cells was observed compared to 20% vehicle treated cells (Figure 2B). MMP-3 levels were significantly reduced following treatment with 10% spruce resin extract, 20% spruce resin extract, and 20% pinoresinol (Figure 2D). The cytokine effects were also compared to appropriate vehicle control using one-way ANOVA with Dunnett’s post hoc test. This resulted in only one change from the previous statistical analysis - IL-1β levels in the 10% pinoresinol treated cells showed a significant increase compared to 10% vehicle treated cells. These experiments showed that the spruce resin extract at 20% exhibits significant anti-inflammatory effects on key pro-inflammatory cytokines related to periodontal inflammation, comparable to that of 0.5 µM dexamethasone. It was also confirmed that the anti-inflammatory effect of the extract was not solely due to pinoresinol. These results suggest that the spruce resin extract at 20% has potential for oral care products targeted at reducing inflammation due to dental plaque.

### 3.4 Anti-Plaque Efficacy of 20% Spruce Resin Extract Toothpaste Formulation

A 20% spruce resin extract-based toothpaste formulation was developed to study the anti-plaque efficacy compared to different treatments (Virbac C.E.T. Enzymatic Toothpaste and Corsodyl Original Mouthwash). The study was done on pooled human saliva-treated rods by measuring mean post-treatment dry biofilm weights and total aerobic colony counts (TVC) achieved after different treatments (Figure 3).

**Figure 3.**
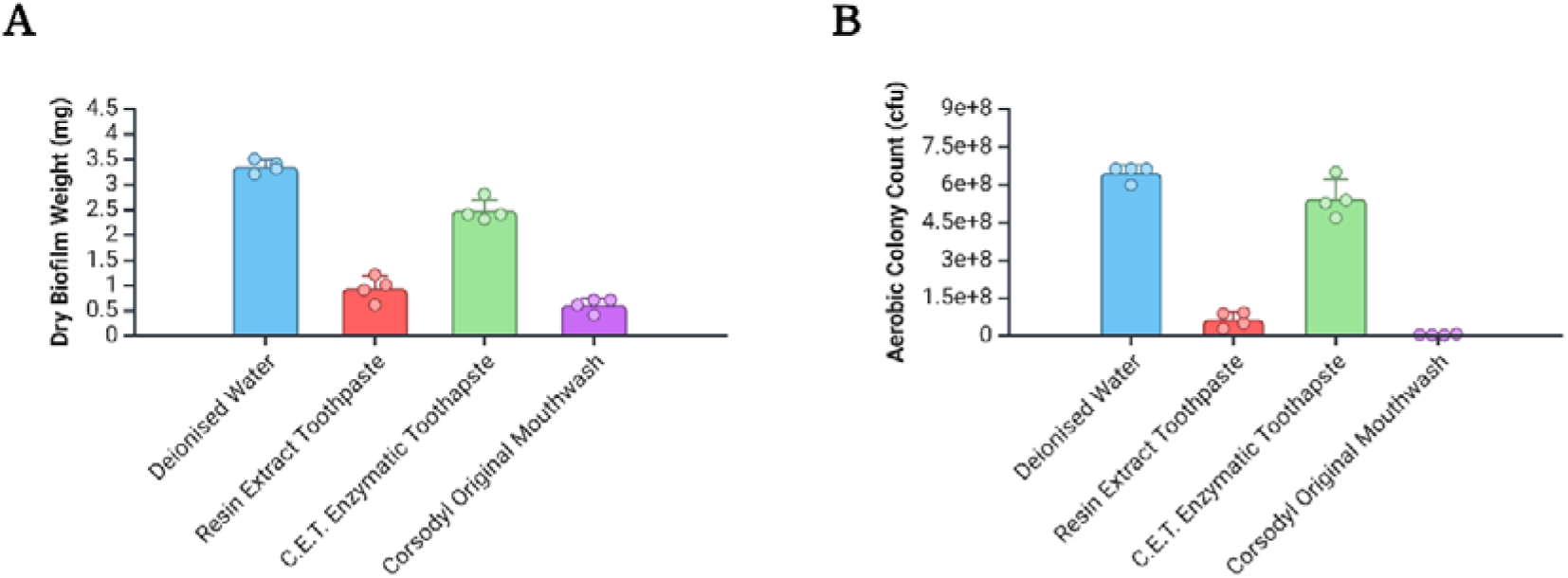
Post-treatment means of (A) dry plaque biofilm weights and (B) total aerobic colony count (TVC).

A Tukey table (Table 3) highlights statistically significant differences between the plaque dry biofilm weights and TVC values achieved by the different treatments.

**Table 3.**
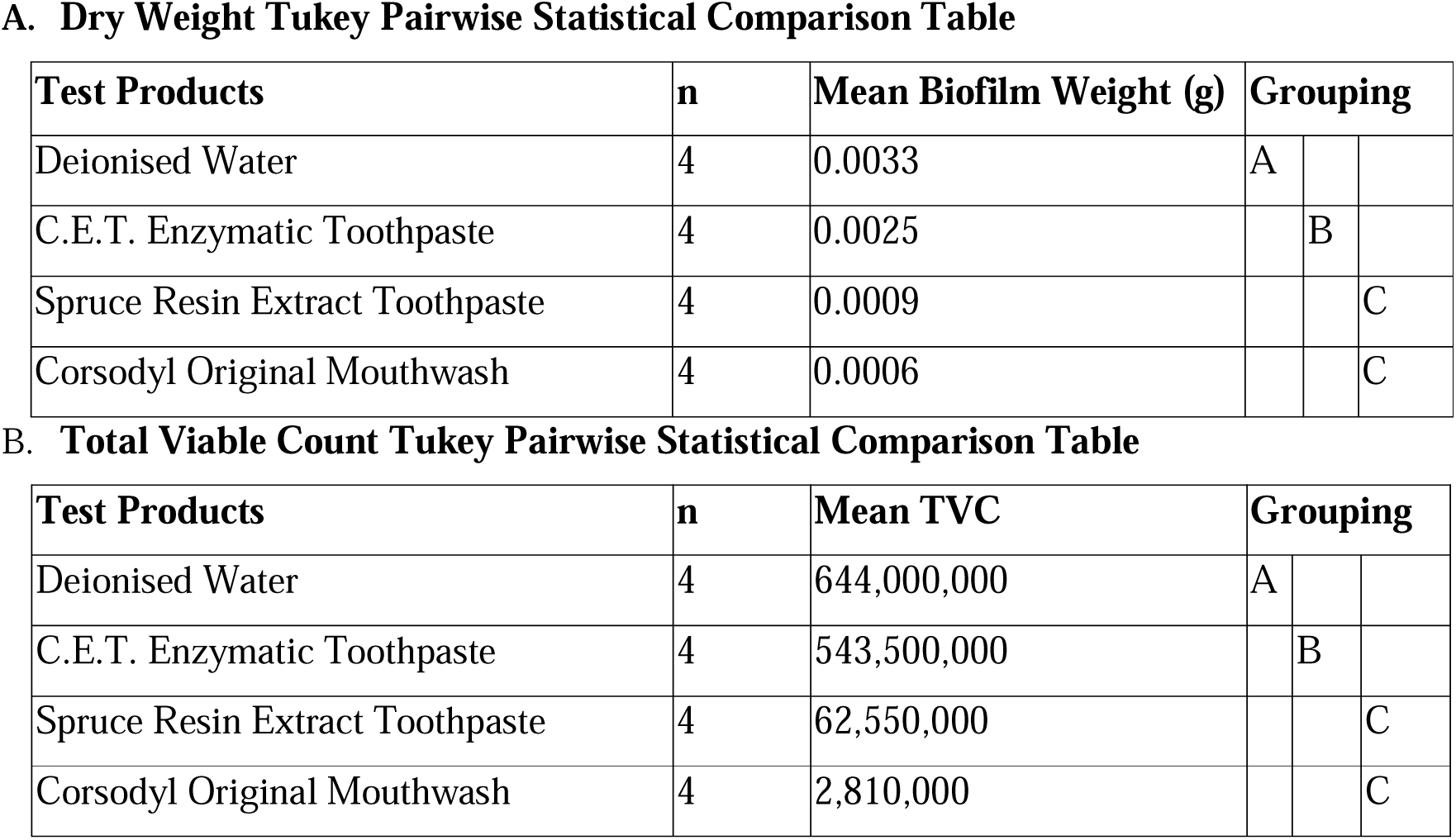
(A) Dry weight and (B) total viable count (TVC). A General Linear Model ANOVA and a Tukey test was used to make pairwise statistical comparisons between the test products and the controls. Means that do not share a letter are significantly different (p ≤ 0.05).

A. Dry Weight Tukey Pairwise Statistical Comparison Table
| Test Products | n | Mean Biofilm Weight (g) | Grouping |  |  |
| --- | --- | --- | --- | --- | --- |
| Deionised Water | 4 | 0.0033 | A |  |  |
| C.E.T. Enzymatic Toothpaste | 4 | 0.0025 |  | B |  |
| Spruce Resin Extract Toothpaste | 4 | 0.0009 |  |  | C |
| Corsodyl Original Mouthwash | 4 | 0.0006 |  |  | C |

B. Total Viable Count Tukey Pairwise Statistical Comparison Table
| Test Products | n | Mean TVC | Grouping |  |  |
| --- | --- | --- | --- | --- | --- |
| Deionised Water | 4 | 644,000,000 | A |  |  |
| C.E.T. Enzymatic Toothpaste | 4 | 543,500,000 |  | B |  |
| Spruce Resin Extract Toothpaste | 4 | 62,550,000 |  |  | C |
| Corsodyl Original Mouthwash | 4 | 2,810,000 |  |  | C |

Corsodyl (containing chlorhexidine digluconate 0.2 % w/v) was selected as a positive control in this test due of its well-established antimicrobial efficacy [36,37]. Sterile water was selected as a negative control since it has no known anti-plaque efficacy. The rods treated with Corsodyl had a significantly (p ≤ 0.05) lower biofilm weight (0.0006g) than the rods treated with sterile water (0.0033g). The total viable count (TVC) data showed also that the rods treated with Corsodyl had a significantly lower (p ≤ 0.05) total viable count (2,810,000 CFU) than the rods treated with sterile water (644,000,000 CFU). The statistically significant differences between the positive control and negative control validated the test’s suitability for comparing the treatments’ effectiveness to inhibit plaque biofilm growth.

Corsodyl treatment led to the lowest mean plaque dry biofilm weight and lowest mean TVC value. The weight and TVC value of Corsodyl was significantly lower than that of the rods treated with the C.E.T. Enzymatic toothpaste (an enzymatic toothpaste with antimicrobial activity [38,39]) slurry (0.0025g and 543,500,000 CFU) and directionally lower than the weight and TVC value from rods treated with the spruce resin extract toothpaste slurry (0.0009g and 62,550,000 CFU). The 20% spruce resin extract toothpaste slurry treatment produced the second lowest mean plaque dry biofilm weight and TVC value. The weight and the TVC value of the spruce resin extract toothpaste were significantly lower than that of rods treated with the C.E.T. Enzymatic toothpaste slurry and the sterile water. The third lowest mean plaque dry biofilm weight and TVC value was observed for the rods treated with the C.E.T. Enzymatic toothpaste slurry. However, the values were significantly lower than that of rods treated with sterile water. Finally, the highest mean plaque dry biofilm weight and TVC value were observed for the rods treated with sterile water. These values were significantly higher than the weights from all other test products assessed in this study.

### 3.5 Proportions of Streptococci

The proportion of streptococci within the total viable counts for the rods treated with the different treatments is shown in Table 4. Streptococci are known to play a key role in dental plaque formation [40,41]. Reducing the prevalence of these microorganisms within the total aerobic colony counts indicates an anti-plaque effect. The identification of streptococci colonies on blood agar plates was primarily based on the type of hemolysis observed around the colonies [42]. The Corsodyl positive control achieved the lowest proportions of streptococci within the total aerobic counts. The 20% spruce resin extract toothpaste slurry achieved the second lowest proportions of streptococci within the total aerobic counts. The C.E.T. Enzymatic toothpaste slurry and sterile water achieved the largest proportions of streptococci with the total aerobic counts.

**Table 4.**
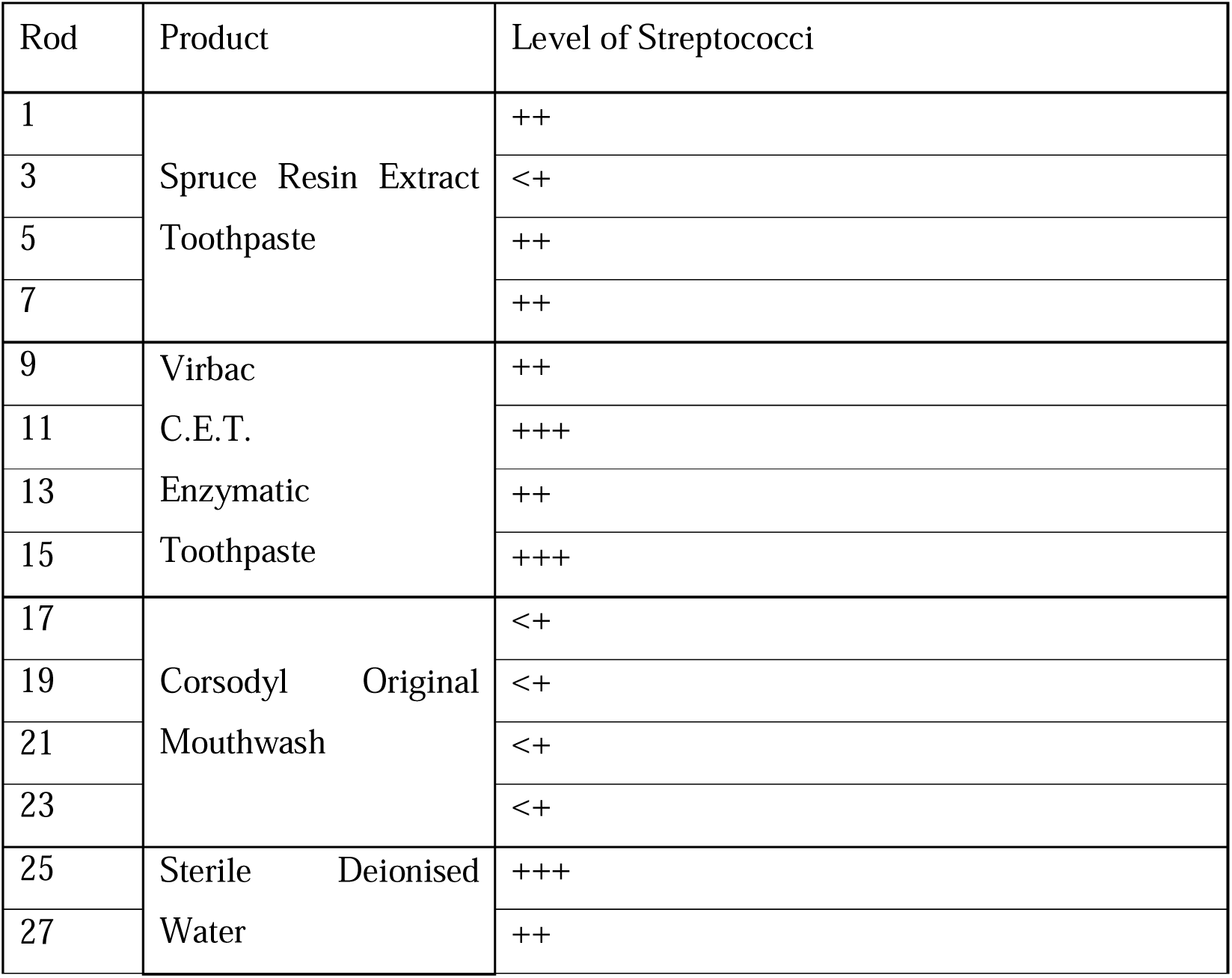
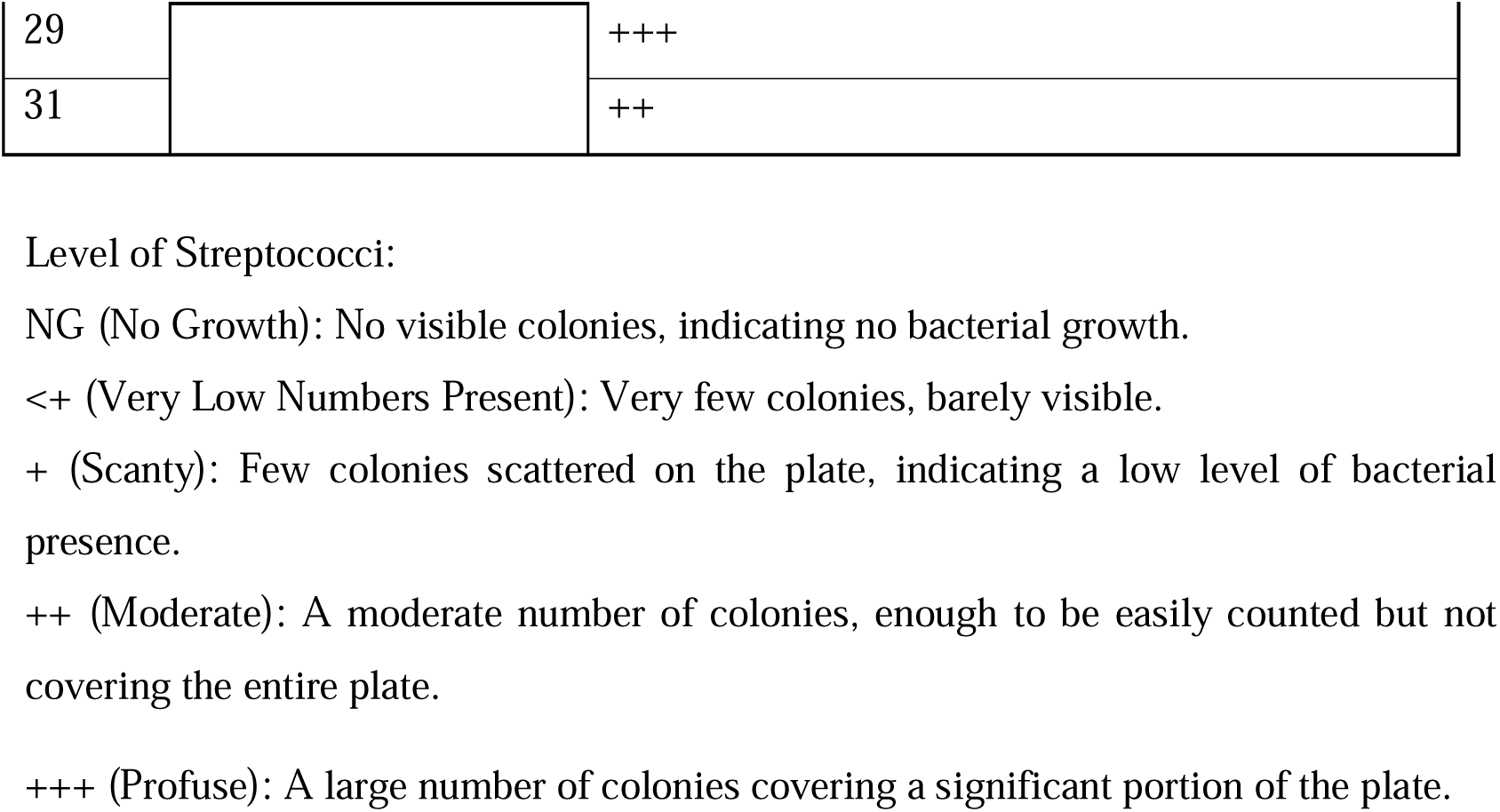
Post-treatment proportions of streptococci with the aerobic counts (TVC).

### 3.6 Oral and Gingival Epithelium Corrosion Tests of Spruce Resin Extract Toothpaste

#### 3.6.1 Measurement of Oral and Gingival Epithelium Cell Vitality (MTT-test), 3 min Incubation

Vitality data of chemicals after an incubation time of 3 min on the oral or gingival epithelium inserts which fall below the vitality data of the non-corrosive negative control, PBS solution, by more than 50 % lead to the evaluation of the substance as oral or gingival epithelium corrosive.

The vitality data (Figure 4A) of the oral epithelium corrosive positive control (glacial acetic acid) of 12.90±1.48 % falls within the valid range of considerably lower than 50 % vitality compared to the PBS negative control for corrosive chemicals after 3 min incubation. The spruce resin extract toothpaste (76.86 ± 11.09 %) did not show an oral epithelium corrosive effect after the 3 min incubation time.

**Figure 4.**
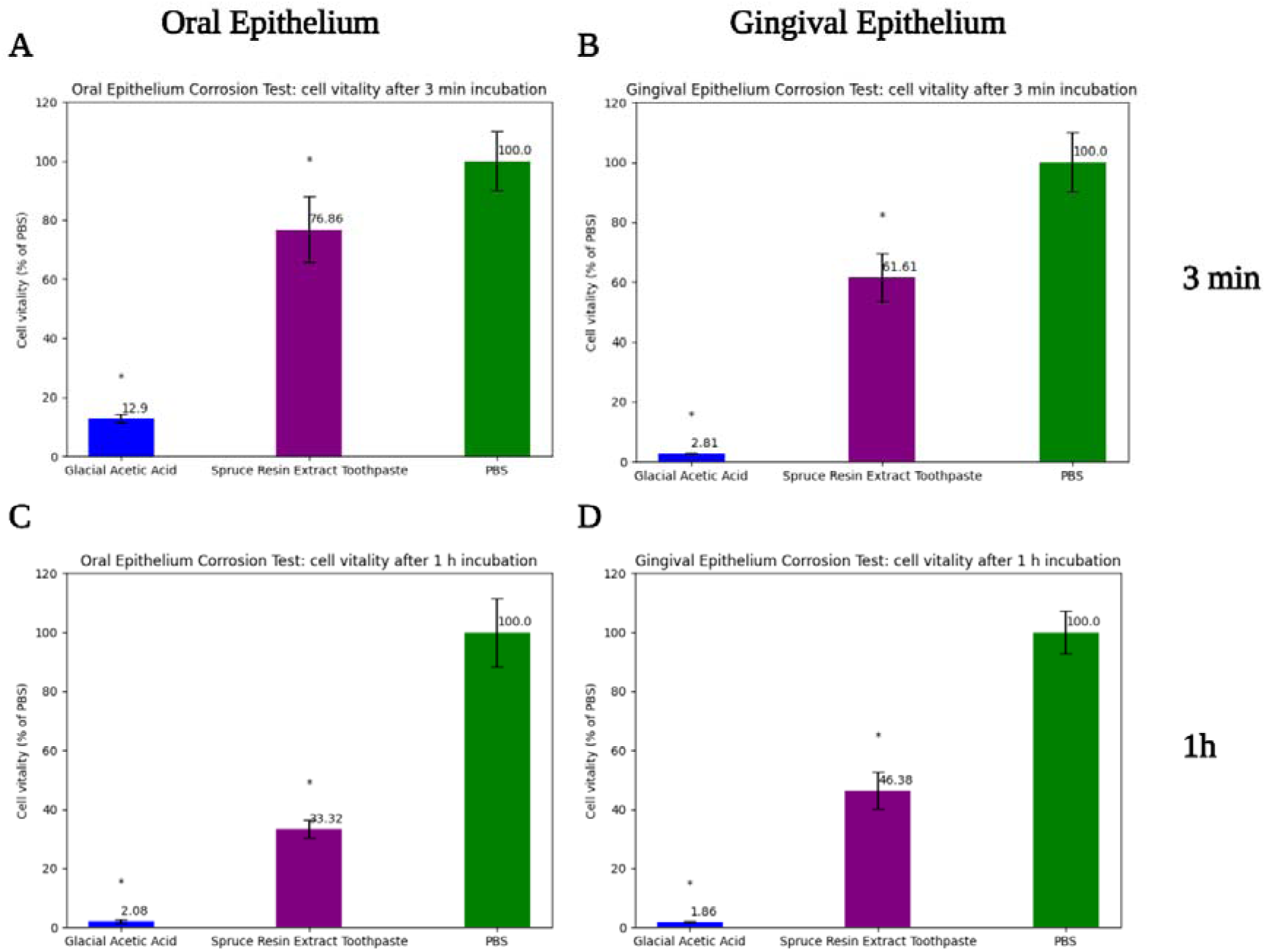
Measurement of cell vitality (MTT-test) of oral epithelium (A and C) and gingival epithelium (B and D) after exposure to the spruce resin extract toothpaste. After 3 min (A and B) and 1h (C and D) incubation times. Positive control - glacial acetic acid, negative control - phosphate buffered saline (PBS). * p < 0.05 Glacial acetic acid vs Spruce resin extract toothpaste.

The vitality data (Figure 4B) of the gingival epithelium corrosive positive control (glacial acetic acid) of 2.81±0.27 % falls within the valid range of considerably lower than 50 % vitality compared to the PBS negative control for corrosive chemicals after 3 min incubation. The spruce resin extract toothpaste (61.61±8.08 %) did not show a gingival epithelium corrosive effect after the 3 min incubation time.

#### 3.6.2 Measurement of Oral and Gingival Epithelium Cell Vitality (MTT-test), 1 h Incubation

Vitality data of chemicals after an incubation time of 1 h on the oral or gingival epithelium inserts which fall below the vitality data of the non-corrosive negative control, PBS solution, by more than 85 % lead to the evaluation of the substance as oral or gingival epithelium corrosive.

The vitality data (Figure 4C) of the oral epithelium corrosive positive control (glacial acetic acid) of 2.08 % falls within the valid range of considerably lower than 15 % vitality compared to the PBS negative control for corrosive chemicals after 1 h incubation. The spruce resin extract toothpaste (33.32±3.07 %) did not show an oral epithelium corrosive effect after the 1 h incubation time.

The vitality data (Figure 4D) of the gingival epithelium corrosive positive control (glacial acetic acid) of 1.86 % falls within the valid range of considerably lower than 15 % vitality compared to the PBS negative control for corrosive chemicals after 1 h incubation. The spruce resin extract toothpaste (46.38± 6.21%) did not show a gingival epithelium corrosive effect after the 1 h incubation time.

The 20% spruce resin extract toothpaste did not cause oral or gingival epithelium corrosive effect. According to OECD 431-TG [43] the oral or gingival epithelium inserts incubated with the substance to be tested need to show at least a cell vitality of 50 % after 3 min incubation and at least a cell vitality of 15 % after 1 h incubation compared to the negative control. These requirements were met after both tested incubation times for oral and gingival epithelium.

## 4. Discussion

In this study, the U937 cells were selected since they differentiate into macrophage-like cells. The macrophages have an important role in inflammation and response to infections [44]. The U937 cells were used to study the production of inflammatory cytokines IL-1β, TNF-α [45], and MMP-3[12] in response to spruce resin extract or pinoresinol at different concentrations. Elevated levels of IL-1β [9,10] and TNF-α are present in periodontitis [46–48]. High levels of IL-1β increase the production of other pro-inflammatory mediators and enzymes, leading to increased inflammation and bone destruction [9,10]. TNF-α also plays an important role in maintaining oral health. TNF-α stimulates proteolytic enzymes that are responsible for the degradation of ECM and bone loss, and increased levels of TNF-α lead to tissue degradation [10,49]. MMP-3 on the other hand is an enzyme that also takes part in ECM degradation by degrading proteoglycans and collagens. It has been found that the MMP-3 might have a destructive effect on gingival tissue in chronic periodontitis [12,50]. MMP-3 is regulated by pro-inflammatory cytokines including those mentioned above (IL-1β and TNF-α). Thus, addressing the reduction of bacterial load, that causes the inflammatory response, is important for the effective management of periodontal disease. The spruce resin extract and pinoresinol (a lignan found at high concentrations in the spruce resin extract, see Table 2) are known to have anti-inflammatory effects [31]. In this study, their potential for reducing gum inflammation was evaluated. It was found that the spruce resin extract at 20% concentration had anti-inflammatory effects on IL-1β, TNF-α, and MMP-3 that are like those of dexamethasone, a potent anti-inflammatory corticosteroid [51]. Reduction of the IL-1β, TNF-α, and MMP-3 levels, by the 20% spruce resin extract could potentially help to prevent gum tissue and bone breakdown, and thus support in the reduction of progression of the periodontal disease. This makes spruce resin extract at this concentration a good candidate for oral health products such as toothpaste for supportive periodontal care.

In this study, a toothpaste formulation containing 20% spruce resin extract was developed and tested for its the non-mechanical anti-plaque efficacy. In addition to 20% spruce resin extract, the toothpaste contained proteolytic and saccharolytic enzymes, and hydrated silica for mechanical plaque removal. The spruce resin [25,26,52,53] and its extracts [20,28,54] are known to possess wide antimicrobial activity also against streptococci [53]. Amyloglucosidase enzyme was selected to break down staches into glucose. Reduction of fermentable carbohydrates available for bacteria could help lowering the risk of dental caries development [55]. The proteolytic enzymes were added due to their promising benefits in dental care [56]. Papain (a papaya fruit derived proteolytic enzyme) was added to help with breaking down protein remains. It also helps in the removal of plaque and promotes the gum health through its anti-inflammatory properties [57]. Bromelain (a pineapple-derived proteolytic enzyme) is also anti-inflammatory and has plaque removal capabilities, adding to the overall effect of the toothpaste [58]. The proteolytic enzymes aid in removing biofilms and pigments *in vitro*, such as theaflavin from dephosphorylated bovine β-casein [59]. Based on clinical studies they can also significantly reduce plaque and gingivitis in orthodontic patients [60]. In addition, they have been found to improve tooth whiteness and are more effective at removing extrinsic stains than conventional toothpastes [56]. This suggests that they may offer benefits over those toothpastes that are only abrasive [61–63]. Moreover, the addition of enzymes in the 20% spruce resin extract toothpaste can assist natural salivary defenses. This is potentially a clinically effective strategy for shifting the balance of the oral microbiome toward improved oral health [3,39].

It was found that the 20% spruce resin toothpaste combined with enzymes was highly effective in reducing plaque biofilm weight and bacterial counts. The toothpaste was almost as effective as a well-established antimicrobial mouthwash (Corsodyl [37]). The substantial reduction in streptococci proportions after toothpaste treatment indicated the effectiveness of the toothpaste formulation containing 20% spruce resin extract in preventing dental plaque formation, where streptococci play a crucial role [40,41,64,65]. The biocompatibility and safety of the spruce resin extract were evaluated on the viability of U937 cells. It was found that most of the treatment’s concentrations studied did not significantly harm the viability of LPS-stimulated U937 cells, except for the 20% vehicle, which reduced viability slightly. Interestingly, pinoresinol at 20% concentration increased the cell viability compared to the vehicle control, suggesting a potential protective effect or enhancement of cell proliferation. However, the spruce resin extract alone, which contains pinoresinol among other compounds, did not show this effect. Since the spruce resin extract at 20% had the highest anti-inflammatory effect on the U937 cells, this concentration was subsequently selected for the toothpaste. The toothpaste’s effect on human 3D-oral and gingival epithelium cell models was also tested to evaluate oral and gingival epithelium corrosion potential of the 20 % spruce resin extract toothpaste formulation combined with enzymes. It was found that the toothpaste did not cause corrosive effect on oral or gingival epithelium. Thus, it was concluded from the biocompatibility studies that 20% spruce resin extract toothpaste lacks significant cytotoxicity at the tested concentrations and is non-corrosive.

## 5. Conclusions

The spruce resin extract at 20% concentration was found to significantly inhibit pro-inflammatory cytokines IL-1β, TNF-α, and MMP-3 in LPS-stimulated U937 macrophage-like cells. The effect was comparable to 0.5 µM dexamethasone. In addition, the toothpaste prepared with 20% spruce resin extract and enzymes reduced dental plaque biofilm weight and total viable bacterial counts at efficacy matching Corsodyl mouthwash. The effect was probably due to antimicrobial compounds found in spruce resin extract such as p-coumaric acid and lignans (e.g. pinoresinol). The toothpaste was also found to be biocompatible for oral use since it was not corrosive on oral and gingival epithelium 3D cell models. These findings also provide scientific validation for the historical use of resins in oral health. By formulating an optimized version of spruce resin extract, it was possible to develop a modern type of natural oral care product that can reduce inflammation and prevent plaque formation. This dual action addresses both major aspects of periodontal disease pathogenesis. To translate these *in vitro* results into real-world practice further research and clinical studies are needed. The spruce resin extract can be also used for developing other oral care formulations e.g., mouthwashes and gels. This aligns well with the growing interest in organic and natural raw materials.

## Data availability

The data supporting the findings of this study are available from the corresponding authors upon reasonable request.

## Acknowledgements

The authors acknowledge the following institutes and their personnel for their cooperation in performing the laboratory tests. Aalto University Bioeconomy infrastructure is thanked for the rent of analysis equipment (GC-MS). The cytokine assays: MD Biosciences, Glasgow, United Kingdom. L. Davies, G. Thomas, and T. Badrock from Intertek ltd, Cheshire, United Kingdom are thanked for the help with anti-plaque testing. Dr. Dietmar Scheddin from CYTOX, Bayreuth, Germany is thanked for corrosion tests according to OECD TG 431. The studies were sponsored by Repolar Pharmaceuticals Oy. Mari Korsu from Teampac Oy is thanked for preparing the toothpaste samples.

## Author contributions

Kamilla Yamileva: Conceptualization, Investigation, Methodology, Data curation, Formal Analysis, Writing – original draft, Visualization, and Writing – review & editing.

Simone Parrotta: Methodology, Investigation, Data curation, and Writing – review & editing.

Evgen Multia: Conceptualization, Investigation, Methodology, Funding Acquisition, Project Administration, Data curation, Formal Analysis, Writing – original draft, Resources, Supervision, Visualization, and Writing – review & editing.

## Declaration of generative AI and AI-assisted technologies in the writing process

During the preparation of this work the authors used OpenAI’s ChatGPT, DeepL Write, and Grammarly in order to improve the readability and language of the manuscript. After using these tools/services, the authors reviewed and edited the content as needed and take full responsibility for the content of the published article.

